# Gut microbial composition modulates food-specific CD4^+^ T cells in food allergy

**DOI:** 10.1101/2025.10.27.684919

**Authors:** Alexa R Weingarden, Flannery S Dahlberg, Caroline N Broude, Xiandong Meng, Sunit Jain, Allison M Weakley, Ashley V Cabrera, Thamotharampillai Dileepan, Michael A Fischbach, Marc K Jenkins

## Abstract

The growing food allergy epidemic is thought to be related to changing environmental factors, particularly changes in the gut microbiome. While prior work has demonstrated that food allergy can be modulated by gut microbes, little is known about how food allergen-specific CD4^+^ T cells are affected by gut microbial composition. Here, we report that food allergy severity differs between mice obtained from two different specific pathogen-free mouse vendors (Jackson Labs [Jax] and Taconic Biosciences [Tac]). Mice from Tac develop diarrhea and anaphylaxis after fewer allergen exposures than mice from Jax. Using food allergen peptide:MHCII tetramers, we also find that Tac mice have fewer allergen-specific regulatory T cells in the small intestine compared to mice from Jax. In addition, Tac mice have a greater abundance of small intestinal mucosal mast cells and increased intestinal permeability. The increased food allergy severity phenotype is transferable via co-housing, which corresponds to a shift in Jax microbial communities towards those found in Tac mice. Our findings demonstrate for the first time that food allergen-specific Treg cells can be modulated by gut microbial community composition, which in turn is correlated to food allergy severity.

## Introduction

The frequency and severity of food allergies have grown rapidly in the past several decades, likely due to changes in our environment^1^. Among other factors, changes to gut microbial composition have been considered a potential cause of this rise in food allergy^2^. If we can understand the interactions between gut microbes and components of the immune system involved in food allergy, we may be able to develop new ways to treat or prevent these allergies.

Prior work on the impact of microbiota in food allergy models is conflicted. In a model of peanut allergy, germ-free mice develop more severe allergic responses, which can be rescued by inoculation with *Clostridia* spores^3,4^. In contrast, in a model of egg allergy, germ-free mice have decreased mast cell homing to the gut and rarely develop symptoms of anaphylaxis^5^. These works have been somewhat limited by the use of germ-free animals with markedly altered mucosal immune systems^6^. Furthermore, no prior work has evaluated whether gut microbial composition impacts the food allergen-specific CD4^+^ T cells which are critical for allergic responses^7^.

Here we report that differences in gut bacterial communities from two groups of specific pathogen-free (SPF) mice drive marked differences in food allergy responses. Animals with more severe allergy responses had a lower proportion of allergen-specific regulatory T cells (Treg) and increased mucosal mast cells (MMCs) in the small intestine and a higher concentration of allergen-specific IgE in serum. Furthermore, co-housing these groups of animals transferred both the more severe allergy phenotype and the microbial community associated with this phenotype. Our findings suggest that gut microbiota can alter food-specific intestinal Treg cell abundance to modulate food allergy.

## Materials and Methods

### Animals

Work in this manuscript was approved by APLAC 32872 from Stanford University and IACUC 2407-42250A from the University of Minnesota. 4-8 week old female BALB/c mice were purchased from either Jackson Labs (#000651) or Taconic Biosciences (#BALB-F). Animals were kept on a standard chow diet for the duration of the experiment. Allergic diarrhea was induced using previous methods^8,9^. In brief, at 6-8 weeks of age, animals were immunized with 50 ug ovalbumin (OVA) (A5503, Sigma-Aldrich; St. Louis, MO, USA) with 1 mg of alum (786-1215, G Biosciences; St. Louis, MO, USA) diluted in 100 ul PBS via intraperitoneal injection. Two weeks following injection, animals were fasted for 3-4 hours during the beginning of the light cycle prior to gavage with 50 mg of OVA (A5503, Sigma-Aldrich) in 300 ul of PBS. Immediately following gavage, animals were placed into individual cages without food but with free access to water. After 60 min, animal cages were observed for diarrhea including loose stools, mucus, and rectal stool staining. Gavage was repeated three times per week for 3-7 total gavages.

### Temperature measurement

1-2 days prior to initial OVA gavage, animals were injected with temperature transponders behind the neck (TP-500, Avidity Science BMDS; Waterford, WI, USA). Baseline temperatures were measured immediately prior to OVA gavage, then every 10 min after gavage up to 60 min, using an electronic reader (DAS-8037-IUS, Avidity Science BMDS).

### Lymphocyte isolation

Lymphocytes from small intestine lamina propria (SI LP) were isolated as previously described^10^. In brief, after removal of mesentery and Peyer’s patches, small intestine was incubated with media containing dithioerythritol (233152, Calbiochem; San Diego, CA, USA) and EDTA (5 mM) to remove the epithelium. The remaining tissue was digested in media containing collagenase I (4197, Worthington, Lakewood, NJ, USA) and DNAse (D5025, Sigma-Aldrich). Lymphocytes were isolated from digested tissue using a Percoll gradient with collection of cells from the interface of 67% and 44% Percoll layers. Lymphocytes from secondary lymphoid organs (lymph nodes and spleen, SLO) were isolated via mechanical dissociation followed by filtration through a 70 um filter to obtain a single-cell suspension.

### Lymphocyte staining

Following isolation, cells were stained with OVA3 peptide:I-A^b^ MHCII tetramers for 1 hour at 37°C in complete cell culture media^11^. Cells isolated from SLO were magnetically enriched following tetramer staining^12^. Next, cells were stained for 30 minutes at 4°C with GhostDye Red 780 (13-0865; Cytek Biosciences, Fremont, CA, USA) and fluorophore-conjugated antibodies specific for: CD4, B220, CD11b, CD11c, F4/80, NK1.1, CD44, ST2 (Table S1). Stained cells were fixed and permeabilized with the FOXP3/Transcription Factor Staining Buffer Kit (00-5523-00; eBioscience, San Diego, CA, USA) according to the manufacturer’s instructions and stained for one hour at room temperature with fluorophore-conjugated Foxp3 and GATA3 antibodies (Table S1). For staining mucosal mast cells, lymphocytes were stained for 30 minutes at 4°C with GhostDye Red 780 and fluorophore-conjugated antibodies specific for: CD4, B220, CD11b, CD11c, Gr-1, CD335, CD3e, CD45, CD117, and ST2 (Table S1). Cells were counted and analyzed by flow cytometry with counting beads on a Cytek Aurora 5-laser flow cytometer or a BD FACSymphony flow cytometer. Data were analyzed using FlowJo software.

### Anti-OVA IgE ELISA

Serum was collected from animals two weeks following IP sensitization and at the time of gut tissue harvest after three OVA gavages. OVA (5 ug/mL) was bound to 96 well plates overnight at 4°C. After washing and blocking with bovine serum albumin (BSA) containing buffer, serum was loaded into wells at four dilutions (1:25, 1:50, 1:100, 1:500) in duplicate. As a standard, mouse anti-OVA IgE (7029, Chondrex; Woodinville, WA, USA) was loaded at eight dilutions (0-12.5 ng/mL) in duplicate in each plate. After incubation and washing, plates were incubated with goat anti-mouse IgE antibody conjugated to horseradish peroxidase (HRP) (1110-05, Southern Biotech; Birmingham, AL, USA). HRP was activated with TMB (555214; BD, Franklin Lakes, NJ, USA) and stopped with sulfuric acid stop reagent (423001; Biolegend, San Diego, CA, USA). Plate absorption was read at 450 nm (with correction at 540 nm) using a GloMax Explorer plate reader (Promega; Madison, WI, USA).

### Intestinal permeability assay

FITC-dextran assay was performed as previously described^13^. In brief, mice were fasted for 4-6 hours. After fast, blood was obtained via facial vein puncture. Next, mice were gavaged with 150 ul of 80 mg/mL 4 kDa FITC-dextran (FD4; Sigma-Aldrich). Facial vein puncture was repeated 4 hours after gavage. Blood was diluted in 15% acid-citrate-dextrose (C3821; Sigma-Aldrich) and centrifuged to remove cells. The resulting plasma was diluted 1:10 and fluorescent signal was excited at 485 nm with detection at 530 nm using Promega GloMax Explorer plate reader. Fluorescence for each sample was normalized to baseline (pre-gavage) fluorescence.

### Microbiome analysis

Genomic DNA was extracted using the DNeasy 96 PowerSoil Pro kit (47014; Qiagen, Venlo, the Netherlands) and quantified in 96-well format using the Quant-iT PicoGreen dsDNA Assay kit (P11496; Thermo Fisher, Waltham, MA, USA). A minimum of 15ng of DNA was taken forward to construct metagenomics sequencing libraries using the Illumina DNA Prep kit, with half-volumes being utilized at each step to minimize cost. 5 cycles of PCR were utilized to amplify DNA and introduce unique dual indices. Post PCR, libraries were purified using a 0.8x bead clean, and were quantified using the Qubit 4 (Thermo Fisher) or the Quant-iT PicoGreen kit. Equal masses of each metagenomics library were pooled, and a dual-sided AMPure XP (A63880; Beckman Coulter, Indianapolis, IN, USA) bead clean was performed on the pooled material to achieve buffer removal and proper size-selection for sequencer loading. The final library pool was quality-checked for size distribution and concentration using the Tapestation 2200 (Agilent Technologies, Santa Clara, CA, USA) and qPCR (BioRad, Hercules, CA, USA). Sequencing was performed on the NovaSeq6000 or NextSeq500 (Illumina, San Diego, CA, USA) using a 2×150bp read configuration, with 20 million paired-end reads being targeted per sample. Raw reads were processed using NinjaMap (an algorithm introduced in Cheng et al., 2022^14^).

Following initial read processing, the microbial composition of the samples was characterized using MetaPhlAn4^15^, a marker gene-based profiling tool that identifies microbial taxa from metagenomic sequencing data. Taxonomic relative abundances were computed based on uniquely mapped marker genes, enabling high-resolution profiling of bacterial communities. Second, microbial community patterns were further analyzed using hclust2^15^, a hierarchical clustering and visualization tool. This method facilitated the generation of heatmaps, providing insights into sample clustering based on microbial composition. Pairwise distances between samples were computed using Bray-Curtis dissimilarity. Lastly, the results were visualized using heatmaps and dendrograms to illustrate microbial abundance patterns and statistical associations.

### Quantification and statistical analysis

Statistical tests were performed using Prism software. The exact values of n (number of animals) are indicated in the figure legends. Outliers were identified by ROUT analysis and removed. p values, derived from unpaired t tests or one-way ANOVA with a Tukey’s multiple comparisons test, are indicated on the figures.

## Results

### Mice harboring distinct gut microbial communities differ in susceptibility to food allergy

We sought to understand whether gut microbial composition affects susceptibility to food allergy. We obtained BALB/c mice from two vendors (Jackson Labs [Jax] and Taconic Biosciences [Tac]) with known differences in gut microbial community composition^16,17^. Mice were sensitized to the model food allergen ovalbumin (OVA) via intraperitoneal injection of OVA with alum as an adjuvant^8^. Two weeks later, animals were challenged with OVA via oral gavage three times weekly for a total of 3-7 gavages (Fig. 1A). Following each OVA challenge, animals were observed for development of diarrhea (loose stool, mucus, and/or anal fecal staining) within 60 minutes following challenge. Consistent with prior findings in this model using mice from Jax^9^, about 50% of Jax animals developed diarrhea after 4-5 gavages (Fig. 1B). However, mice from Tac developed diarrhea substantially faster, with 74% of animals developing diarrhea after only 3 gavages (Fig. 1B).

**Figure 1.**
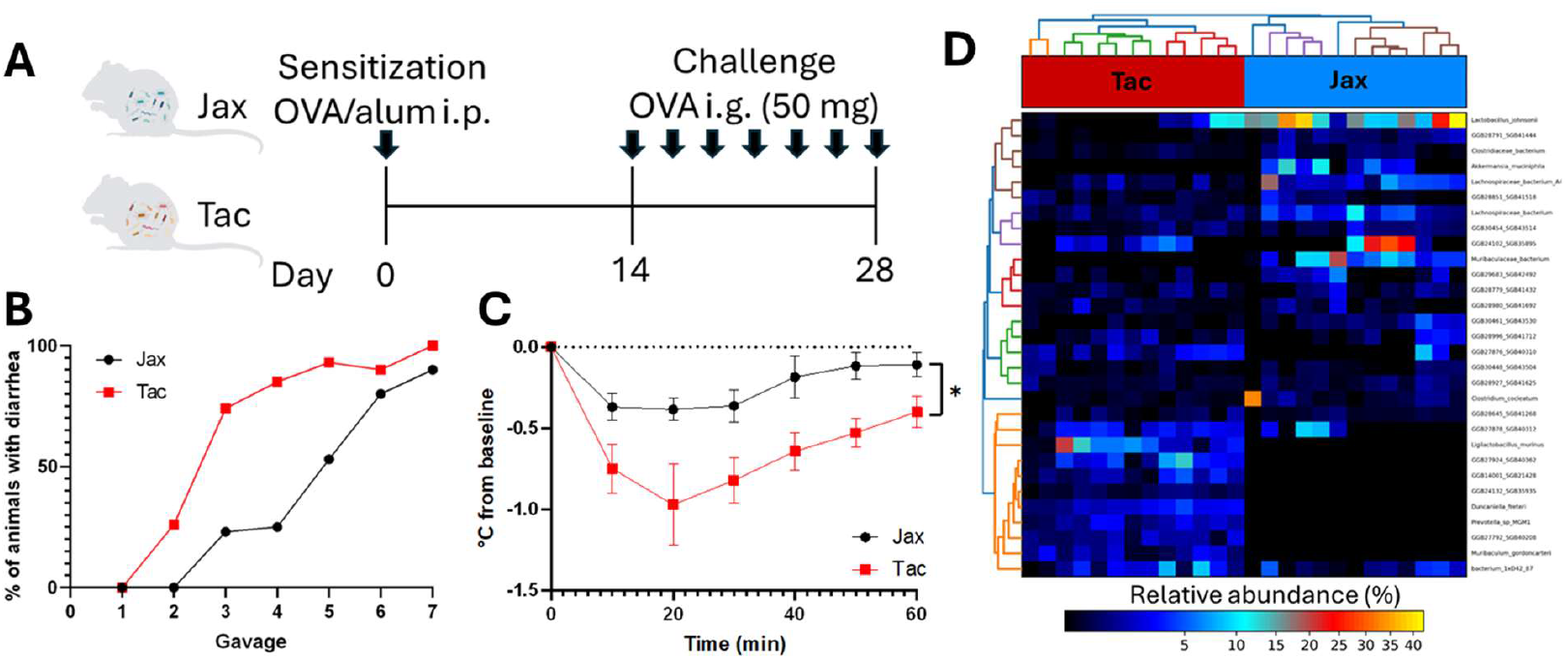
Severity of allergic diarrhea is associated with differences in fecal bacterial communities. A) Disease model schematic. Animals were sensitized to OVA via intraperitoneal (i.p.) injection with alum on day 0. Beginning on day 14 mice received intragastric (i.g.) challenge with OVA every 2-3 days, for up to 7 challenges. B) Percentage of animals from Jackson Labs (Jax) and Taconic Biosciences (Tac) with diarrhea following each OVA gavage (n = 10-35 per time point per group). C) Body temperature change from baseline within 60 min following the third OVA challenge gavage (n = 8-10 per group). *p < 0.05. D) Heatmap of amplicon sequence variants (ASVs) with significantly different relative abundance in stool from Tac and Jax mice.

We next sought to examine if these differences in allergen-driven diarrhea corresponded to differences in anaphylactic shock. As a proxy for shock, we tracked body temperature of mice before and up to 60 minutes after OVA challenge. Following the third OVA gavage, animals from Tac became significantly more hypothermic than Jax animals, suggesting that anaphylactic shock was more severe in Tac mice (Fig. 1C).

We then confirmed that gut microbial communities were distinct in animals from Jax and Tac. We performed metagenomic sequencing analysis of stool pellets from mice from Jax and Tac and found that communities from animals from each vendor clustered more closely to each other than animals from different vendors (Fig. 1D). These distinctions were driven by several amplicon sequence variants (ASVs), including increased abundance of *Lactobacillus johnsonii, Akkermansia muciniphila*, and *Clostridium cocleatum* in Jax mice and increased abundance of *L. murinus, Duncaniella freteri, Prevotella* sp. MGM1, and *Muribaculum gordoncarteri* in Tac mice. These findings suggest that stool microbial composition in mice correlates with differences in allergy severity.

### Mice with greater food allergy susceptibility have lower abundance of allergen-specific Treg cells in small intestine

Anaphylaxis in food allergy is dependent on allergen-specific Th2 cells^7^. We therefore investigated whether differences in OVA-specific CD4^+^ T cells could explain the accelerated anaphylaxis seen in Tac mice using an OVA peptide (OVA3):I-A^d^ MHCII tetramer. In secondary lymphoid organs (SLO; spleen and lymph nodes), sensitization with OVA and alum followed by three oral OVA challenges expanded the OVA3:I-Ad tetramer-binding CD4^+^ population by 9 to 29-fold compared to naïve animals (Fig. 2A, Fig. S1A). The total number of tetramer-positive cells was similar in both sensitized Jax and Tac animals (Fig. 2A). Although tetramer-positive cells were predominantly CD44^hi^ in allergic mice (Fig. 2B), indicating antigen experience, very few cells in animals from either vendor expressed either Foxp3 or markers of the Th2 lineage (ST2 and GATA3^18^) (Fig. 2C-D).

**Figure 2.**
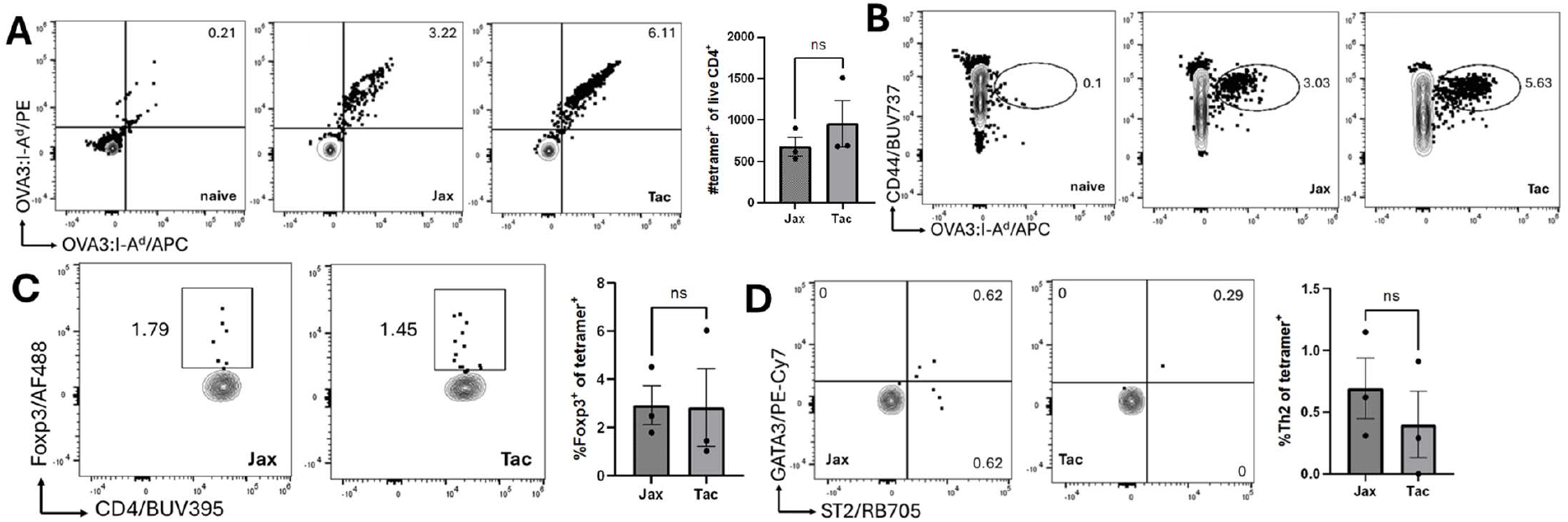
Allergen-specific CD4^+^ T cells expand but express few Treg and Th2 cell markers in SLO. A) Dual tetramer staining for OVA-specific CD4^+^ T cells in SLO (n = 2-3 per group). B) Expanded OVA3:I-A^d^ tetramer binding CD4^+^ cells are CD44^hi^. C) Tetramer^+^ Treg cells in Jax and Tac mice. D) GATA3^+^ST2^+^ Th2 tetramer^+^ cells in Jax and Tac mice (gated on Foxp3^-^). ns, not significant.

We next explored whether the phenotype of OVA-specific cells differed at the site of allergen challenge, the SI LP. As expected in non-lymphoid tissue, tetramer-positive CD4^+^ T cells were virtually undetectable in naïve animals but represented 0.2-2.95% of CD4^+^ cells in sensitized and orally challenged mice (Fig. 3A, Fig. S1B). After three challenges, there were significantly more OVA-specific cells in Tac compared to Jax SI LP (Fig. 3A). Notably, a significantly lower proportion of tetramer-positive cells in Tac animals were Treg cells (Fig. 3B), as were non-allergen specific Treg cells in SI LP (Fig. S2A). Of the remaining tetramer-positive cells in animals from both vendors, the majority (20-62.5%) co-expressed ST2 and GATA3, suggesting a Th2 phenotype (Fig. 3C). However, the abundance of tetramer-positive and tetramer-negative Th2 cells was not significantly different between Tac and Jax mice (Fig. 3C, Fig. S2B).

**Figure 3.**
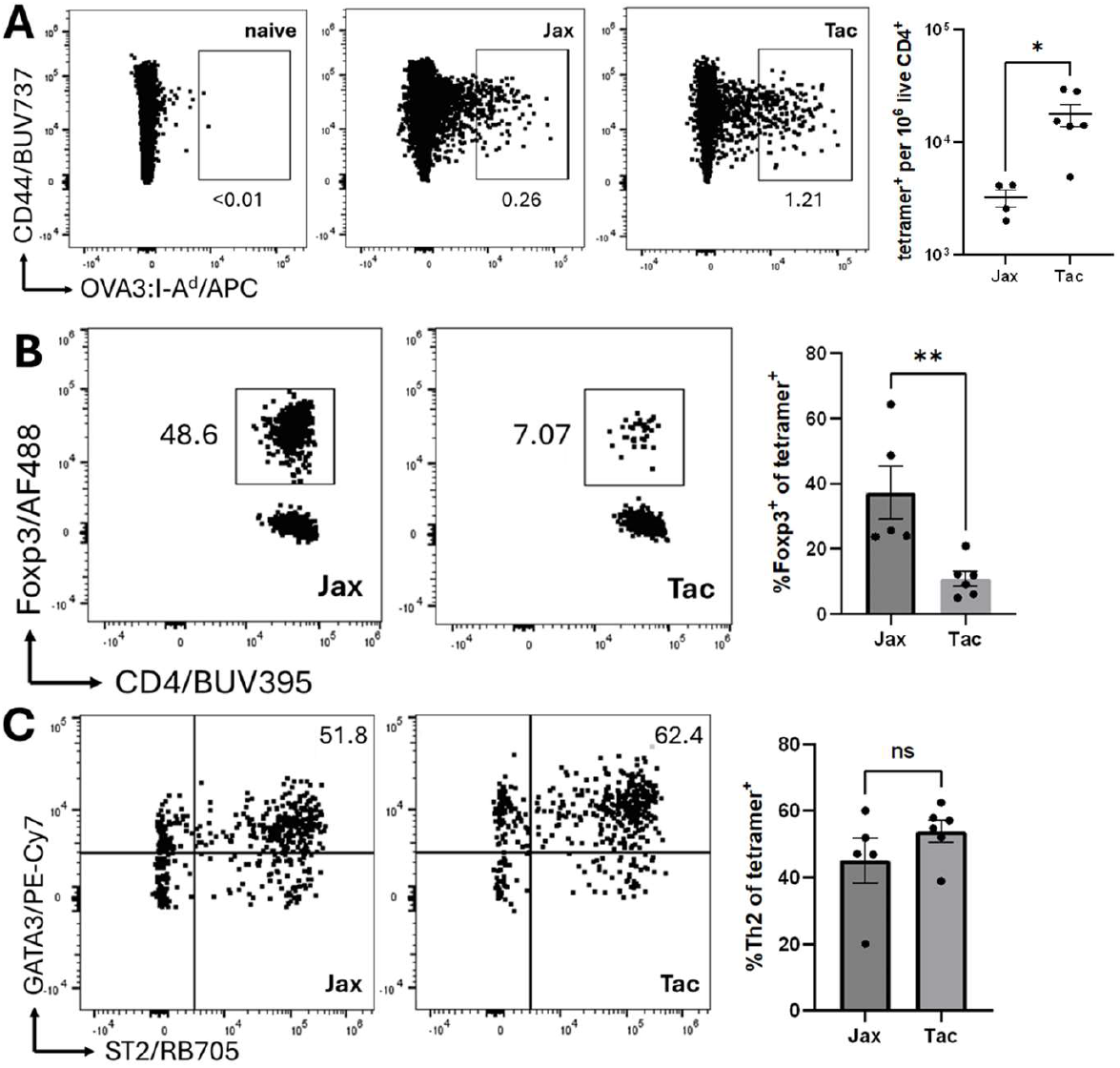
OVA-specific Treg cells are decreased in Tac SI LP lymphocytes. A) OVA3:I-A^d^ tetramer-positive CD4^+^ T cells in SI LP from naïve (left) versus sensitized and triple oral challenged Jax (middle left) and Tac (middle right) animals. Proportion of tetramer-positive cells in Jax and Tac animals per 10^6^ live CD4^+^ T cells (right) (n = 4-6 per group). B) Proportion of Foxp3^+^ Treg cells within tetramer-positive population (n = 5-6 per group). C) Proportion of tetramer-positive ST2^+^GATA3^+^ Th2 cells within the Foxp3^-^ population in (B) (n = 5-6 per group). *p < 0.05; **p < 0.01; ns, not significant.

These findings suggest that gut microbial composition can drive an alteration in food allergen-specific Treg cell abundance in SI LP, which in turn is correlated to allergy severity.

### Mice with worsened food allergy severity have higher abundance of mucosal mast cells and increased intestinal permeability

We next sought to understand how a reduced proportion of allergen-specific Treg cells in SI LP contribute to allergy. OVA-specific IgE is required for both diarrhea and anaphylaxis in this model^19^. In other models, the serum concentration of allergen-specific IgE has correlated with the severity of anaphylaxis as measured by hypothermia following allergen challenge^3,4,20^. Although OVA-specific IgE was detectable in the sera of animals following sensitization, but the concentration was not significantly different in Jax or Tac animals (Fig. 4A). However, following three oral challenges with OVA, serum anti-OVA IgE was significantly higher in animals from Taconic (Fig. 4A).

**Figure 4.**
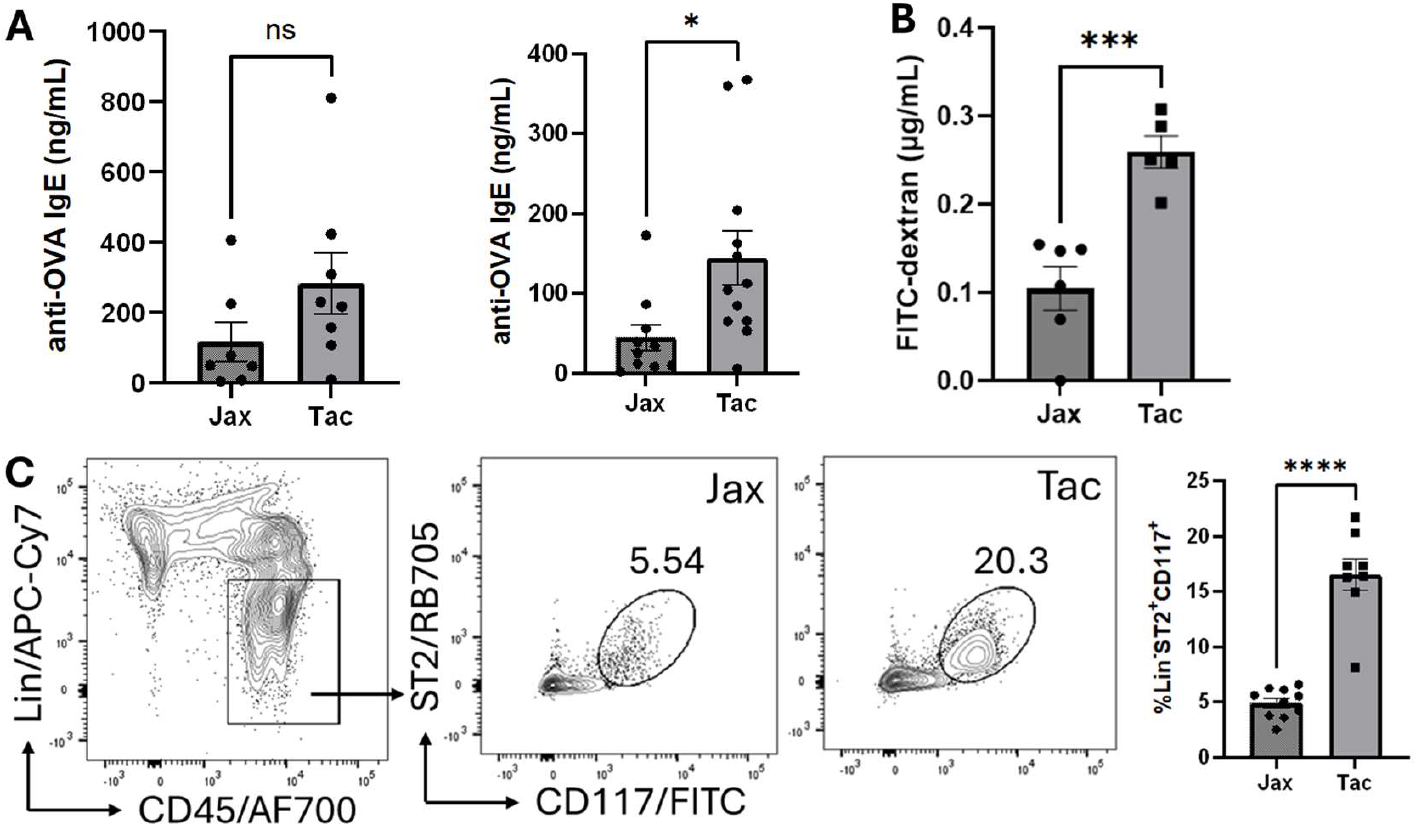
Anti-OVA IgE, gut permeability, and SI LP mucosal mast cell abundance are higher in Tac animals. A) Serum anti-OVA IgE concentration after 2 weeks post-sensitization (left) and after 3 OVA gavage challenges (right) (n = 7-12). B) Concentration of FITC-dextran in serum following gavage in naïve 6-9 week old Jax or Tac animals (n = 5-6 per group). C) Abundance of Lin^-^ST2^+^CD117^+^ mucosal mast cells in SI LP after 3 OVA gavages (n = 8-10 per group). *p < 0.05; ***p < 0.001; ****p < 0.0001; ns, not significant.

Recent work has demonstrated that increased uptake of intestinal allergen is associated with increased MMC degranulation and food allergy severity^21^. We therefore examined differences in intestinal permeability in Jax and Tac mice, and indeed found that uptake of FITC-dextran was increased in Tac mice (Fig. 4B).

Mucosal mast cells (MMCs) are the critical final effectors of anaphylaxis in this and other models of food allergy^8,9^. In addition to promoting B cell class switching to IgE, Th2 cells are also thought to initiate MMC migration to and expansion in SI LP, vial IL-9 production^9^. In contrast, Treg cells can inhibit mast cell proliferation and degranulation^22–25^. Therefore, we investigated whether differences in abundance of OVA-specific Treg cells were correlated to differences in SI LP MMC abundance. Indeed, the abundance of CD117^+^ST2^+^ MMCs^9^ (Fig. S3) was significantly higher in Tac mice compared to Jax mice after the three oral OVA challenges (Fig. 4C).

Overall, these findings suggest that mice from different animal vendors have differences in allergen-specific Treg cells in SI LP, serum allergen-specific IgE, MMC abundance, and intestinal permeability, in association with differences in allergy severity.

### Food allergy susceptibility is transmissible by early-life transfer of microbial communities

Next, we sought to determine if allergy severity was indeed modulated by gut microbial composition. To do so, we co-housed Jax and Tac mice for four weeks beginning shortly after weaning before OVA priming and oral challenge (Fig. 5A). In contrast to Jax animals housed only with mice from the same vendor, Jax mice co-housed with mice from Tac developed allergen-driven diarrhea at the same accelerated rate as mice originally from Tac (Fig. 5B). Notably, this worsening of food allergy severity was age-dependent, as food allergy severity did not transfer when animals were co-housed beginning at age six weeks (Fig. S4).

**Figure 5.**
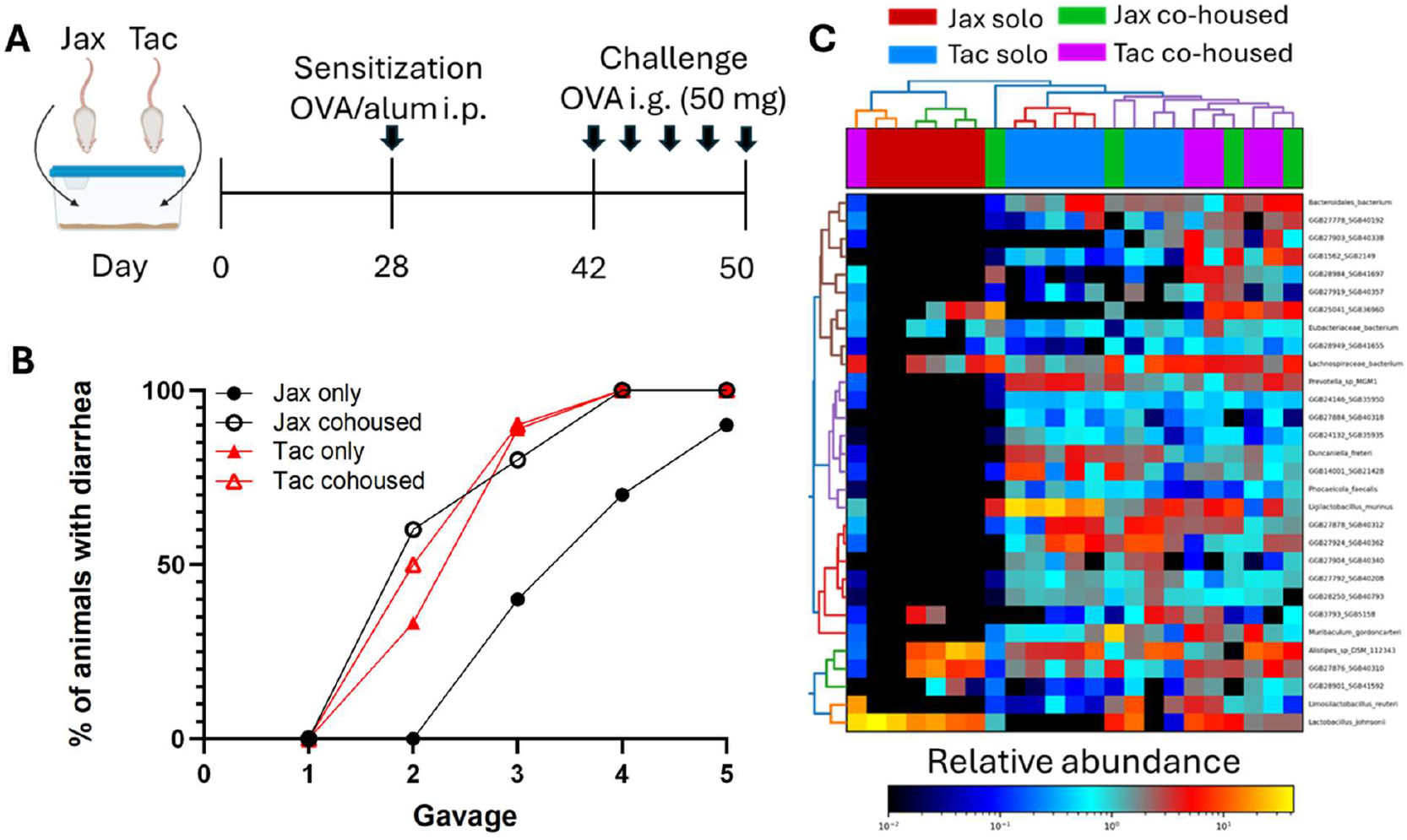
Food allergy susceptibility is transferable via co-housing. A) Mice from Jax and Tac were co-housed for 28 days prior to OVA sensitization and subsequent OVA gavages. B) Percentage of cohoused versus non-cohoused (only) animals from Jax and Tac with diarrhea following each OVA challenge (n = 10 per group). C) Heatmap of ASVs with significantly different relative abundance in stool from Tac and Jax co-housed and non-cohoused (solo) mice.

To investigate whether the change in allergy severity corresponded to a change in gut microbial community, we performed shotgun metagenomic sequencing of co-housed mice. Indeed, mice originally from Jax that had been co-housed with Tac mice acquired fecal bacterial communities that more closely resembled those from Tac mice than animals from Jax (Fig. 5C). Community differences were driven by an increase in *L. murinus, D. freteri, Prevotella* sp. MGM1, *M. gordoncarteri, Limosilactobacillus reuteri*, and *Phocaeicolia fecalis* and a decrease in *L. johnsonii* in co-housed Jax mice. These findings suggest that shifts in fecal bacterial communities could drive the changes we observed in allergy severity.

All together, our findings suggest that differences in gut microbial composition affect allergen-specific Treg abundance and intestinal permeability, as well as the concentration of allergen-specific IgE following oral challenge, which in turn can affect the abundance of mucosal mast cells. These findings could explain why mice with a distinct microbial composition have worsened food allergy severity.

## Discussion

A major goal of this research was to characterize allergen-specific CD4^+^ T cells in food allergy and determine if their behavior is affected by differences in gut microbial composition. Most investigation of food allergen-specific T cells has been performed using adoptively transferred cells expressing a transgenic TCR such as DO11.10 cells^23^ or via *ex vivo* stimulation of cells with allergen^26^. We present here a characterization of *in vivo* native food allergen-specific T cells using MHCII tetramer staining. This revealed the surprising finding that although they are antigen-experienced (CD44^hi^), allergen-specific cells in secondary lymphoid organs do not have clear lineage markers for either Th2 or Treg cells, even when mice have detectable allergen-specific IgE in serum and display anaphylaxis upon oral allergen challenge. Instead, allergen-specific cells have a marked Th2 (and Treg) phenotype within peripheral tissue at the site of allergen challenge, in the small intestinal lamina propria. Notably, mice harboring distinct gut microbial populations develop a different abundance of allergen-specific Treg cells in the SI LP, which negatively correlates with allergy severity.

A key remaining question is whether the increased population of OVA-specific Treg cells in SI LP is directly responsible for allergy protection seen in Jax mice. It has been shown that adoptive transfer of Treg cells, including allergen-specific DO11.10 Treg cells generated *in vitro*, can inhibit anaphylaxis^23,27^. However, DO11.10 Treg cells generated from an allergy-prone host failed to provide this protection^23^. Of note, because these cells were generated *in vitro* from transgenic animals, these results may not accurately reflect *in vivo* conditions in animals possessing a full native set of TCRs^28^. Furthermore, a highly non-physiological number of DO11.10 Treg cells were transferred, which may have altered the phenotype or function of the transferred cells^28^. Future work could investigate whether transfer of tetramer-isolated allergen-specific Treg cells we describe here can protect from anaphylaxis.

Future work should also focus on the interaction between mast cells and antigen-specific Treg cells in food allergy. Prior work has suggested that both Treg cells and mast cells can suppress the other. For example, Treg cells can inhibit mast cell degranulation *in vitro*, a process thought to be mediated by OX40-OX40L interaction^22^. Conversely, mast cells may suppress the generation of Treg cells, through an unclear mechanism involving mast cell Syk signaling^24^. Whether increased Treg cells in Jax mice suppress the proliferation or degranulation of mast cells, or if increased mast cells in Tac mice suppress Treg cells, is currently unclear.

Our work expands on existing findings regarding the role of gut microbial composition in food allergy. In two different models of food allergy, germ-free mice exhibit evidence of increased allergic responses, including worsened hypothermia upon allergen challenge and increased serum concentration of allergen-specific IgE^3,4^. These changes were reversed when mice were colonized with *Clostridiales* species, but not with *Proteobacteria* nor in mice with genetic ablation of the transcription factor Rorγt in Treg cells. However, it is unclear whether these microbial clades impacted allergen-specific T cells, or if they drove a bystander effect via induction of microbe-specific Treg cells. Our work clearly demonstrates that allergen-specific Treg cells are influenced by gut microbial composition.

Gut barrier function has previously been shown to be a potential mechanism by which gut microbes can modulate food allergy. One group has found that increases in the short-chain fatty acid (SCFA) butyrate produced by *Clostridia* species can decrease intestinal permeability to allergen, which is associated with decreased anaphylaxis^3,29^. Another group recently demonstrated that indole metabolites produced by several bacterial species are also protective against food allergy^30^. While gut barrier function was not examined in that work, these indole metabolites can decrease intestinal permeability^31,32^. In addition, recent work has shown a clear link between increased intestinal allergen uptake via secretory epithelial cells and food allergy susceptibility^21^. Our findings are in line with these existing data and support the concept that modulation of intestinal allergen absorption is a key mechanism whereby gut bacteria can influence the outcome of food allergy. Furthermore, our findings suggest that differences in gut antigen permeability may impact food allergen-specific T cell phenotype. In contrast to this prior work, however, we found that different bacterial taxa were associated with worsened food allergy. In fact, mice with worsened food allergy from Tac had significantly increased stool abundance of ASVs corresponding to *L. murinus* and *L. reuteri*, which produce protective indole metabolites^30^. Although ASVs corresponding to *Clostridium cocleatum*, a butyrate-producing species, were more abundant in Jax mice, this taxon was not significantly different in abundance in Jax mice co-housed with Tac animals (Fig. 5C). Additional work analyzing bacterial metabolites directly may help us understand whether SCFA, indole derivatives, or novel metabolites may drive the differences in allergy phenotype we observed in our study.

Finally, the limitation of allergy phenotype transfer to the peri-weaning period suggests that susceptibility is set during the weaning reaction^33^. This phenomenon describes intestinal immune maturation in response to the influx of novel microbial species during weaning. Mice with altered microbial composition due to antibiotic treatment at the time of weaning fail to induce this reaction, resulting in later susceptibility to Th2 cell-driven colonic inflammation. This weaning reaction also drives the induction of Treg cells in the colon. It is likely that a similar reaction occurs in the small intestine during weaning, and perhaps exposure to different microbial communities during this critical period also drives differences in food allergy phenotype we find here.

We propose a model wherein microbial composition affects gut permeability, in turn affecting food allergen uptake in the intestine (Fig. 6). Increased allergen absorption might limit the differentiation of allergen-specific Treg cells in the small intestine, in turn removing a check on mucosal mast cell abundance and/or degranulation.

**Figure 6.**
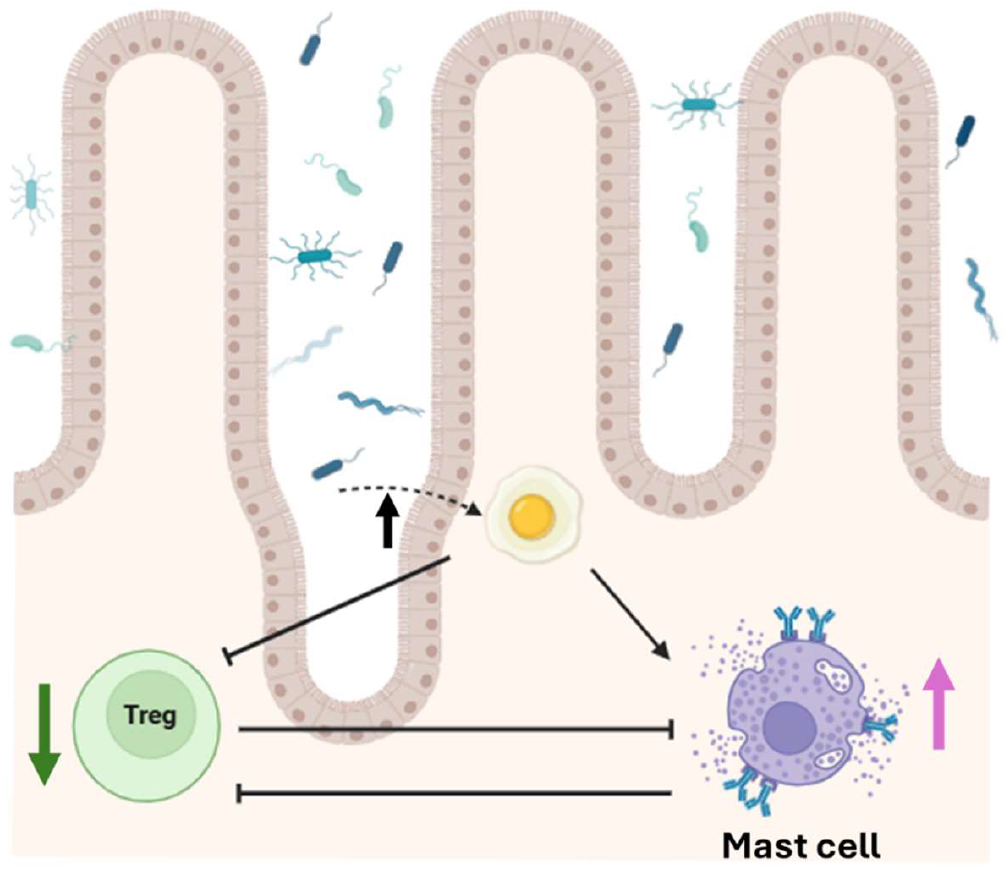
Proposed model for modulation of food allergy severity by gut microbes. Microbial composition in Tac mice promotes increased gut permeability and uptake of food allergen. In turn, this may either limit the development of food-specific Treg cells directly or could promote degranulation and expansion of mucosal mast cells. This increased mast cell population could then suppress intestinal Treg cells. Ultimately, these changes result in increased food allergy severity, particularly an allergic response after fewer exposures of oral allergen.

Alternatively, increased absorption could more directly result in increased serum allergen-specific IgE and mucosal mast cells, which then could inhibit allergen-specific Treg development. Future work investigating microbial and immune mechanisms behind the relationships in this model could deepen our understanding of how microbial composition alters tolerance to food antigens.

## Conclusions

One hypothesis for the rise in food allergies over the past several decades is changes in the gut microbiome. Our findings here support this hypothesis, suggesting that even two different SPF microbial populations can modulate food allergy. Critically, worsened allergy susceptibility was paralleled by a lower population of allergen-specific Treg cells in the small intestine, revealing previously unexplored effects of gut microbes on food-specific Treg cells. In addition, our findings suggest that altered gut permeability may be one mechanism by which microbial communities can influence food-specific Treg cell development. Future work will need to focus on which microbes can enhance gut permeability, and whether increased antigen in the intestine directly limits Treg cell development or supports mast cell expansion, indirectly inhibiting Treg cells in food allergy. Long-term, these results could lead to novel microbial therapeutics for food allergy and could also shed light on other disorders of food antigens such as celiac disease.

## Acknowledgements

The authors wish to thank Dr. Alexander Khoruts for critical manuscript feedback. Flow cytometry analysis for this project was done with the assistance of the University of Minnesota Flow Cytometry Resource Core and the Stanford Shared FACS Facility (RRID: SCR_017788). Some data were collected on an instrument in the Stanford Shared FACS Facility obtained using NIH S10 Shared Instrument Grant (1S10OD026831-01).

## Funding

This research was supported by funding from NIAID (R01AI187164). ARW was supported by the Life Science Research Foundation postdoctoral fellowship and Open Philanthropy. ARW holds the following patent: Compositions and methods for transplantation of colon microbiota, US Patent 2014/0147417 A1, filed March 9, 2012, accepted May 29, 2014. MAF is a cofounder of Kelonia and Revolution Medicines, a member of the scientific advisory boards of the Chan Zuckerberg Initiative, NGM Biopharmaceuticals and TCG Laboratories/Soleil Laboratories, and an innovation partner at The Column Group.

## Author Contributions

ARW designed and performed research and wrote the manuscript; FSD, CNB, XM, SJ, AMW, and AVC performed research; TD provided reagents; and MAF and MKJ designed the research.

---

**Supplemental Figure 1.**
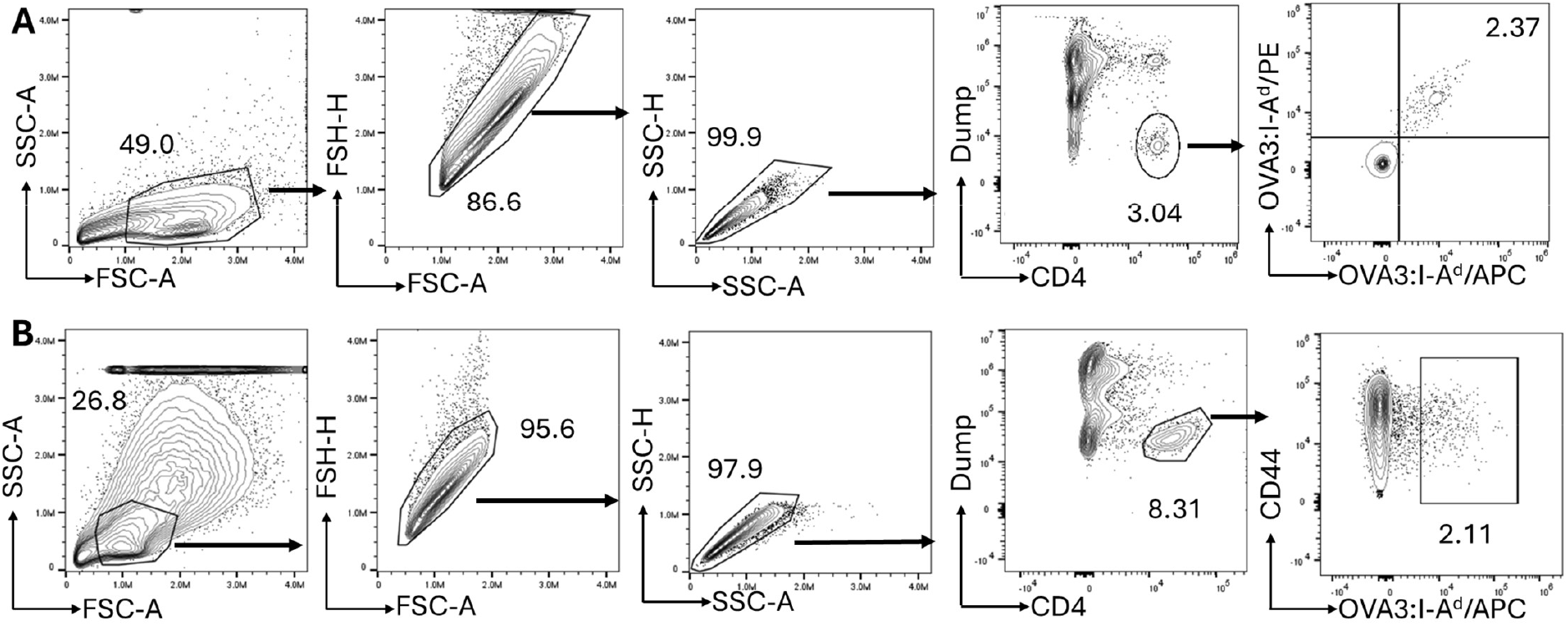
Gating for tetramer-positive spleen and lymph node (SLO) (A) and small intestine lamina propria (SI LP) (B) cells from Balb/c mice sensitized to OVA via OVA-alum IP injection and orally challenged three times with OVA via gavage. SLO cells were magnetically enriched following OVA3:I-A^d^ tetramer staining. This gating was designed to capture cells with light scatter properties of lymphocytes which were singlets, were live cells which did not express non-T lineage markers (Dump; B220, CD11b, CD11c, F4/80, NK1.1), and which were CD4^+^. SI LP dump gating also included staining for TCRγδ.

**Supplemental Figure 2.**
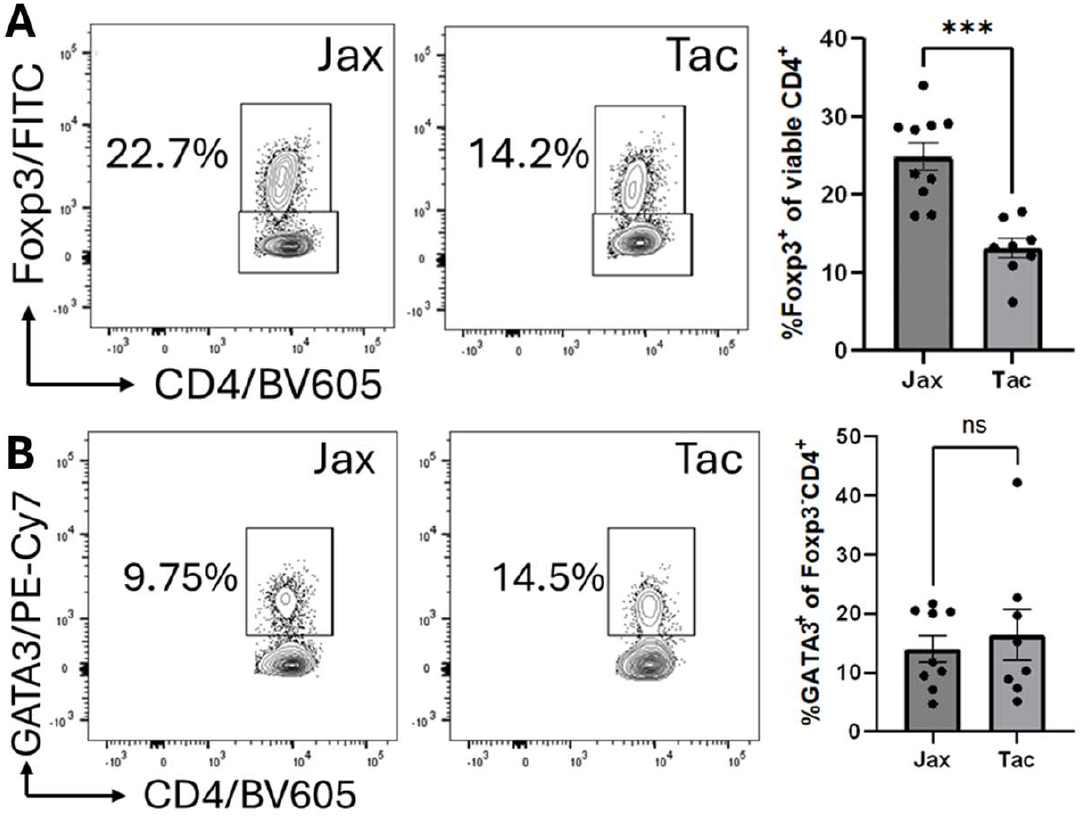
Tetramer negative CD4^+^ T cell populations in Jax and Tac mice in SI LP. A) Proportion of Treg cells among tetramer-negative live CD4^+^ T cells (n = 8-10 per group). B) Proportion of GATA3^+^ cells within the Foxp3^-^ gate in (A) (n = 8-9 per group). ***p < 0.001; ns, not significant.

**Supplemental Figure 3.**
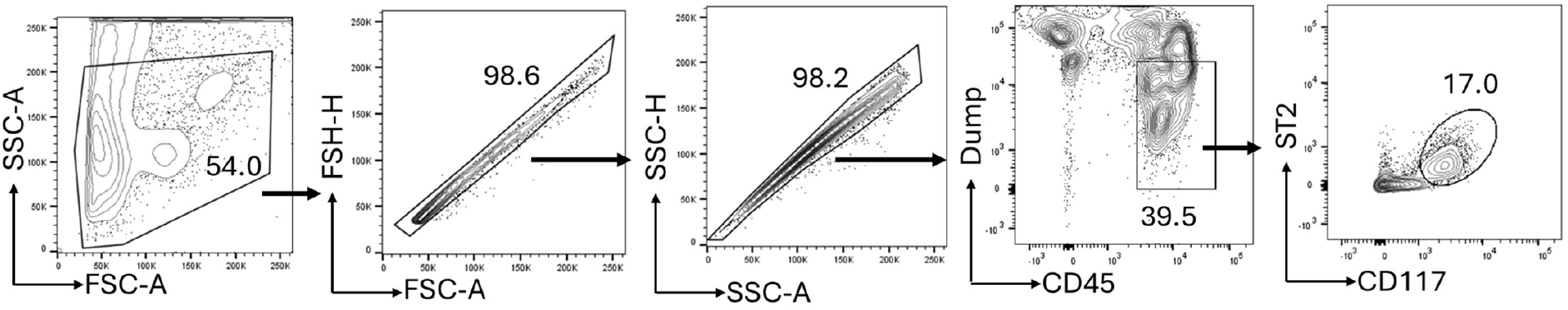
Gating for mucosal mast cells (MMC) from SI LP. This gating was designed to capture cells with light scatter properties of leukocytes which were singlets, were live CD45^+^ cells which did not express lineage markers (Dump; B220, CD11b, CD11c, Gr-1, CD335, CD4, CD3e), and which were CD117^+^ and ST2^+^.

**Supplemental Figure 4.**
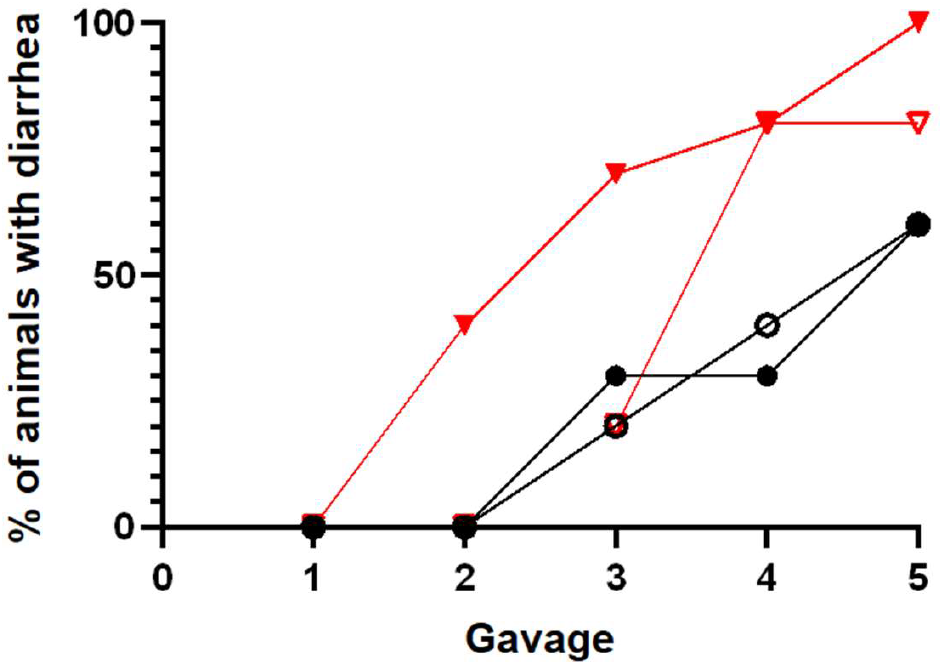
Transfer of allergy phenotype is age-dependent. Percentage of Jax or Tac co-housed or non-cohoused (only) mice with diarrhea after each OVA gavage (n = 5). Animals were co-housed at age 6 weeks for 2 weeks prior to sensitization.

**Table S1.**
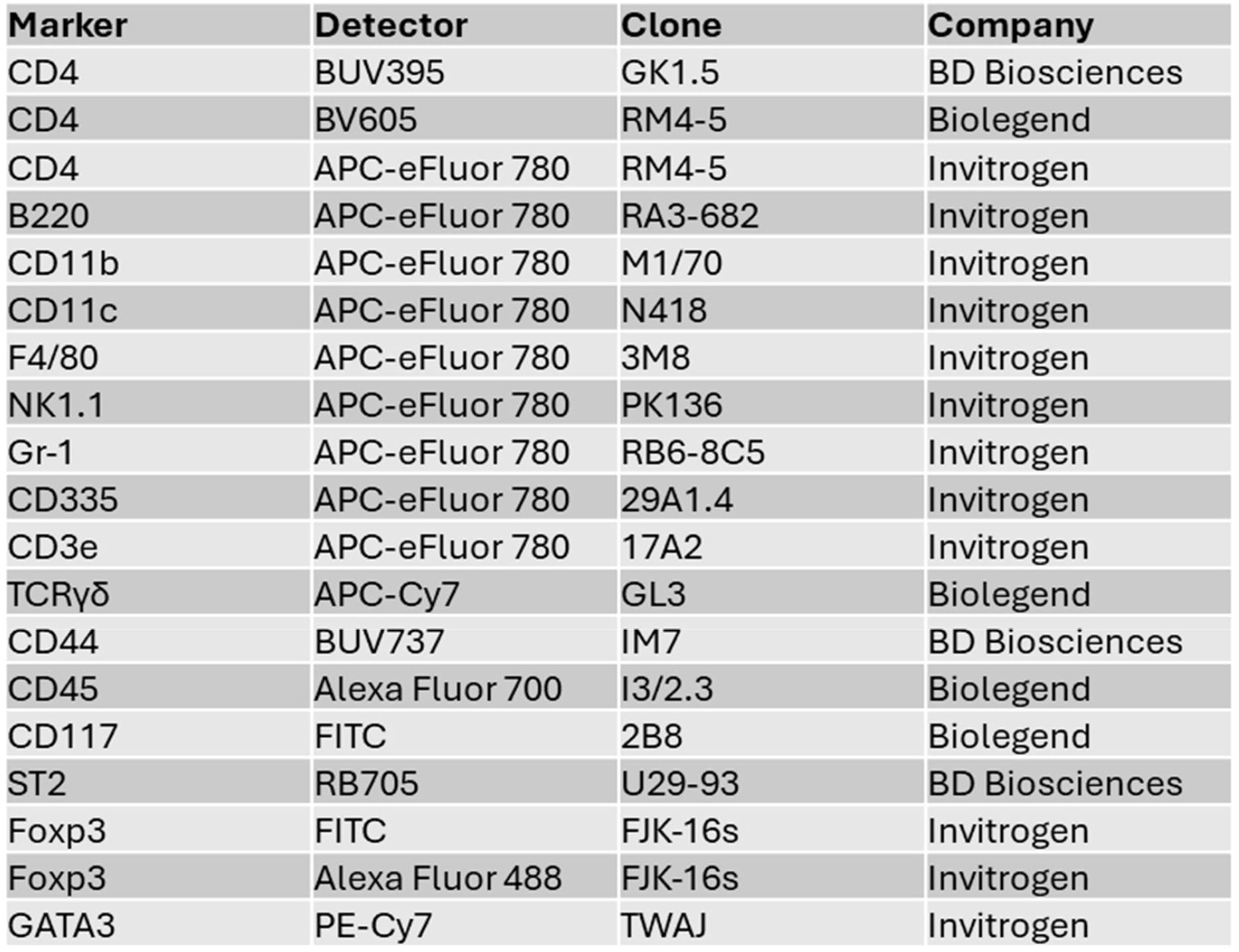
Antibodies used in this study.

## Notes

### Competing Interest Statement

The authors have declared no competing interest.

